# The deletion of aldehyde:ferredoxin oxidoreductase-encoding genes in *Clostridium ljungdahlii* results in changes in the product spectrum with various carbon sources

**DOI:** 10.1101/2024.07.20.604392

**Authors:** Saskia T. Baur, Sarah Schulz, Joshua B. M Cluskey, José Antonio Velázquez Gómez, Largus T. Angenent, Bastian Molitor

**Author notes:** These authors contributed equally to the work (Saskia T. Baur, Sarah Schulz). Corresponding author: Bastian Molitor. Max Planck Institute for Chemical Energy Conversion, Stiftstrasse 34-36, 45470 Mülheim a. d. Ruhr, Germany.

## Abstract

Biofuels, such as ethanol, can be produced by the microbial fermentation of waste gases that contain carbon dioxide (CO_2_) and carbon monoxide (CO). The acetogenic model microbe *Clostridium ljungdahlii* converts those substrates into acetyl-CoA with the Wood-Ljungdahl pathway. During autotrophic conditions, acetyl-CoA can be reduced further to ethanol *via* acetic acid by the enzymes aldehyde:ferredoxin oxidoreductase (AOR) and alcohol dehydrogenase. Here, the genes encoding both tungsten-dependent AORs (*aor1*, CLJU_c20110 and *aor2*, CLJU_c20210) were deleted from the genome of *C. ljungdahlii*. Ethanol formation was enhanced for *C. ljungdahlii* Δ*aor1* with different carbon sources, that is, fructose, a mixture of hydrogen (H_2_) and CO_2_, and CO. The highest and lowest ethanol:acetic acid ratio was detected during growth with H_2_/CO_2_ and CO, respectively. Oscillating patterns were observed during growth with CO, underpinning the importance of a balanced redox metabolism.

## 1. Introduction

It is crucial to reduce greenhouse gas emissions, foremost CO_2_, to stop global warming from exceeding 2°C above pre-industrial temperatures. Implementing carbon capture and storage or utilization technologies can be part of the solution (Baker et al., 2020). One such technology is the production of commodities from waste gases with bacterial (syngas) fermentation (Fackler et al., 2021). Acetogenic bacteria, such as *Clostridium ljungdahlii* and *Clostridium autoethanogenum*, are model microbes for syngas fermentation (Abrini et al., 1994; Fackler et al., 2021; Köpke et al., 2010; Tanner et al., 1993). They utilize the Wood-Ljungdahl pathway, which allows the conversion of H_2_, CO_2_, and CO into the central intermediate acetyl-CoA from which biomass is derived (Fackler et al., 2021).

Acetyl-CoA can be further converted into ethanol *via* the direct or the indirect route, the latter involving the enzyme aldehyde:ferredoxin oxidoreductase (AOR) (Köpke et al., 2010; Liew et al., 2017; Richter et al., 2016). AOR is a key component for autotrophic growth and ethanol production of *C. ljungdahlii* and *C. autoethanogenum* during syngas fermentation (Richter et al., 2016; Valgepea et al., 2017). *C. ljungdahlii* and *C. autoethanogenum* encode three AORs, two of which are tungsten-dependent (AOR1, CLJU_c20110, CLAU_0089; and AOR2, CLJU_c20210, CLAU_0099), and one of which is molybdenum-dependent (AOR3, CLJU_c24130, CLAU_0452) (Köpke et al., 2010; Liew et al., 2017; Whitham et al., 2015). Previously, the *aor1* and *aor2* genes were inactivated for *C. ljungdahlii* by introducing premature stop codons *via* CRISPR-targeted base editing, which resulted in decreased acetic acid titers for both strains in H_2_/CO_2_-fed cultures (**Tab. S1**) (Xia et al., 2020). However, ethanol production was not changed for the *aor1*-inactivated strain, and completely abolished for the *aor2*-inactivated strain with H_2_/CO_2_ (**Tab. S1**). Growth with CO was not investigated (Xia et al., 2020). For *C. autoethanogenum*, the *aor1* and *aor2* genes were completely deleted, and the strains were characterized under different growth conditions (**Tab. S1**) (Liew et al., 2017). When grown with CO, the deletion of *aor1* led to decreased ethanol titers, whereas the deletion of *aor2* led to increased ethanol titers compared to the wild type strain (**Tab. S1**). The authors did not investigate all strains with H_2_/CO_2_ (Liew et al., 2017). Thus, the impact of *aor*-gene deletions on the growth and product profiles of model acetogens is not yet fully understood (**Tab. S1**). The results for *C. autoethanogenum* suggest that AOR1 prefers catalysis toward acetaldehyde, while AOR2 prefers the direction toward acetic acid (Liew et al., 2017). However, for *C. ljungdahlii*, the directionality cannot be deduced from the available data. We hypothesize that the directionality of AORs for *C. ljungdahlii* is the same as for *C. autoethanogenum* because the *aor*-gene sequences are identical between the two microbes. In this study, we constructed *C. ljungdahlii* deletion strains that lack the entire *aor1*- and *aor2*-gene individually and as a double deletion, as opposed to the *aor*-inactivated strains from Xia et al. (2020). We then physiologically characterized all strains with fructose, H_2_/CO_2_, and CO as substrates to further understand the role of AOR for syngas fermentation.

## 2. Materials and Methods

### 2.1 Bacterial strains and growth conditions

All strains, plasmids, and primers used and constructed are listed in **Tables S2-S4**. *Escherichia coli* was used for cloning and methylation of plasmid DNA as described before (Xia et al., 2020). *C. ljungdahlii* was cultivated anaerobically at 37°C with agitation of 150 rpm in <u>R</u>einforced <u>C</u>lostridial <u>M</u>edium (RCM) or PETC medium with 5 g L^-1^ fructose under an atmosphere of N_2_ (Klask et al., 2022). The media were supplemented with clarithromycin (5 μg mL^-1^) and anhydrotetracycline (100 ng mL^-1^) when appropriate (Klask et al., 2022; Klask et al., 2020). Heterotrophic growth experiments were conducted in 250-mL serum bottles (Glasgerätebau Ochs Laborfachhandel e.K., Bovenden, Germany). For autotrophic growth, fructose was omitted from the PETC medium, and the headspace was exchanged by three vacuum/gassing cycles to H_2_/CO_2_ (80/20 vol-%) or CO (100 vol-%) at 1 bar overpressure. The growth experiments were conducted in 1-L DURAN pressure plus bottles (VWR International, LLC, Darmstadt, Germany), which were inoculated with cultures that had been precultured in 50 mL RCM and then in 100 mL PETC medium with fructose, and concentrated with PETC medium without fructose.

### 2.2 Cloning procedures

Standard molecular cloning and DNA manipulation techniques were applied as described before (Klask et al., 2022; Xia et al., 2020). If not noted otherwise, all enzymes and kits were purchased from New England Biolabs GmbH (Frankfurt, Germany). A functional CRISPR/Cas9-based gene deletion system for *C. ljungdahlii* was implemented. The plasmid pMTLCas9 was constructed by reverting the deactivating point mutations of *dcas9* in pMTLdCas9 (Xia et al., 2020) by PCRs on the inner part of *cas9* and the rest of the backbone, followed by ligation using the NEBuilder HiFi DNA Assembly Master Mix. The plasmids pgRNA_*aor1* and pgRNA_*aor2* were constructed by inverse PCRs on the plasmid pTargetF (Jiang et al., 2015) with appropriate primers and digesting the PCR products with *Dpn*I. These plasmids allowed to amplify the sgRNA constructs targeting the respective *aor* gene. The up- and downstream regions of *aor1* as homology repair templates and the *aor1*-targeting sgRNA were ligated with the linearized pMTLCas9 plasmid (*Sal*I), which yielded pMTLCas9_*aor1*. For constructing the plasmid to delete *aor2*, the linearized pMTLCas9 was ligated with the *aor2*-targeting sgRNA while maintaining the *Sal*I restriction site. The up- and downstream regions of *aor2* as homology repair templates were subcloned into pMiniT2.0 using the NEB PCR Cloning Kit, which yielded the plasmid pMiniT2.0_donor-*aor2*. From there, the homology repair templates were amplified and ligated with the *Sal*I-linearized pMTLCas9-aor2-STEP1, resulting in pMTLCas9_*aor2*.

### 2.3 Genetic engineering of *C. ljungdahlii*

Methylation of the plasmids pMTLCas9_*aor1* or pMTLCas9_*aor2* and transformation of *C. ljungdahlii*, as well as recovery and induction of *cas9*-expression, was performed as described before (Molitor et al., 2016; Xia et al., 2020). Colonies were picked and screened with colony PCR for the presence of the plasmid and deletion of *aor1* and *aor2*. The deletion was verified by Sanger sequencing, and positive clones were cured from the plasmid. The double-deletion strain was generated by implementing an additional *aor2* deletion in the plasmid-cured *C. ljungdahlii* Δ*aor1* strain. After genomic DNA preparation using the MasterPure Gram positive DNA Purification Kit (LucigenCorp., Middleton, WI, USA), the DNA libraries of the final deletion strains were prepared with a Ligation Sequencing Kit V14 according to the manufacturer’s protocol, and sequenced with a Nanopore MinION sequencer (Oxford Nanopore Technologies, Oxford, United Kingdom) (Lu et al., 2016). Genome assembly was performed with the EPI2MELABS wf-bacterial-genome pipeline (Wang et al., 2021), which applied FlyE v2.9.3-b1797 (Kolmogorov et al., 2020). Sequence analysis was performed using Geneious Prime 2024.0.3 (https://www.geneious.com).

### 2.4 Analysis of growth, product formation, and gas consumption

Growth of the *C. ljungdahlii* strains was determined by measuring the optical density with a BioMate 160 (Thermo Fisher Scientific Inc., Waltham, MA, USA) at 600 nm (OD_600_). The pH was measured with a micro pH electrode LE422 (Mettler-Toledo GmbH, Gießen, Germany). Fructose, acetic acid, ethanol, lactate, and 2,3-butanediol were measured from randomized samples by HPLC on a Shimadzu LC20 system (Shimadzu, Kyoto, Japan) with additional filtration of samples (syringe filters, pore size 0.22 μm, Carl Roth GmbH) after thawing (Klask et al., 2020). Gas analysis was performed with gas chromatographs equipped with a Molsieve 13X column or a Haysep D column (SRI GC, Torrance, USA) to detect CO, N_2_, and O_2_, or H_2_ and CO_2_, respectively (Rohbohm et al., 2023).

### 2.5 Statistical analysis of the results of the growth experiments with fructose and H_2_/CO_2_

For statistical analyses, first, the parametric assumptions normality (Shapiro-Wilk) and homogeneity of variances (Levene) were tested for the growth rate and products at the endpoint of the growth experiments. If the assumptions were met, a 1-way ANOVA with a Tukey HSD as a *post-hoc* test was used to test for the difference in growth rate, and in the acetate and ethanol production and their ratio. If the assumptions were not met, a non-parametric Kruskal-Wallis test was used with a Kruskal-Wallis 1-way ANOVA as a *post-hoc* test with *p*-values given as *p*_kw_. IBM SPSS Statistics v.29 software (IBM Corp., Armonk, NY, USA) was used to perform the statistical analyses with the assumptions 0.05 for the α error and 95 % for the significance.

## 3. Results and Discussion

### 3.1 Growth with fructose displays an effect of *aor* deletions on ethanol formation

We constructed the strains *C. ljungdahlii* Δ*aor1*, *C. ljungdahlii* Δ*aor2*, and *C. ljungdahlii* Δ*aor1*Δ*aor2* successfully with a CRISPR/Cas9-based gene deletion system. First, all strains, including *C. ljungdahlii* wild type (wt) as a reference, were cultivated with fructose as a carbon source (**Fig. 1**). All strains grew, but the OD_600_ decreased considerably after reaching the maximum for *C. ljungdahlii* Δ*aor2* (**Fig. 1A**). While this effect was also observed for the other strains, it was much less pronounced (**Fig. 1A**). The pH dropped from 5.82 ± 0.01 to 3.75 ± 0.08 for all strains (**Fig. 1B**). *C. ljungdahlii* Δ*aor2* did not consume the provided fructose completely, while the other strains did (**Fig. 1C**). This observation was accompanied by a lower total maximum acetic acid and ethanol titer for *C. ljungdahlii* Δ*aor2* compared to the other strains (**Fig. 1D,E**). *C. ljungdahlii* Δ*aor1*Δ*aor2* produced a similar amount of acetic acid than *C. ljungdahlii* Δ*aor2* (*p*=0.978), which was significantly lower compared to *C. ljungdahlii* wt (*p*=0.001) and *C. ljungdahlii* Δ*aor1* (*p*=0.014, **Fig. 1D, Tab. S5**). Although the production of ethanol for *C. ljungdahlii* Δ*aor1*Δ*aor2* was delayed compared to the other strains, this strain reached a significantly higher ethanol titer than *C. ljungdahlii* wt, *C. ljungdahlii* Δ*aor2* (*p*=0.001 for both), and *C. ljungdahlii* Δ*aor1* (*p*=0.006) in this experiment (**Fig. 1E, Tab. S5**). At the end of the experiment, *C. ljungdahlii* Δ*aor1*Δ*aor2* and *C. ljungdahlii* Δ*aor1* had similar (*p*=0.665) but significantly higher ethanol:acetic acid (E:A) ratios compared to the other two strains (*p*=0.001 for both, **Fig. 1F, Tab. S5**). The different *aor*-deletion strains were previously constructed in the closely related species *C. autoethanogenum* (Liew et al., 2017). Contrary to our findings, these strains exhibited more pronounced differences in the E:A ratio with fructose (**Tab. S1**,**S5**). *C. autoethanogenum* Δ*aor2* achieved an E:A ratio of 1.2, whereas *C. autoethanogenum* Δ*aor1* and *C. autoethanogenum* Δ*aor1*Δ*aor2* achieved an E:A ratio of ∼0.4, respectively. These ratios are either higher (1.2) or lower (0.4) compared to *C. autoethanogenum* wt, which reached an E:A ratio of 0.75 (Liew et al., 2017). This indicates differences in the physiological role of the AORs for the two species and their preferred catalytic reaction direction, as further discussed below. The deletion of the *aor* genes, in the previous studies and our results, reveals changes in the E:A ratio or growth behavior among the different strains with fructose. Thus, the AORs also seem to play a role during heterotrophic growth (Richter et al., 2016).

**Fig. 1.**
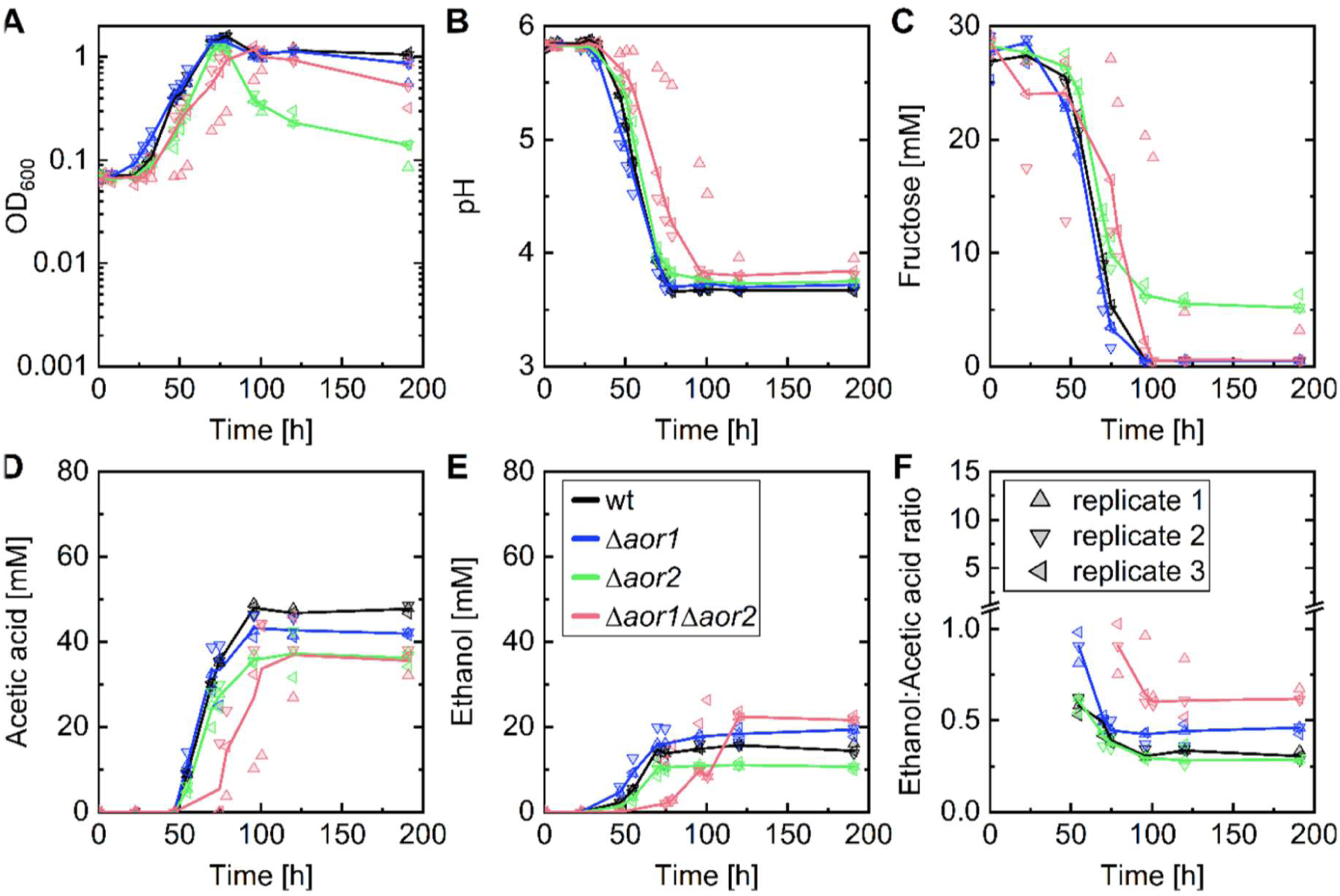
Cultivation of *C. ljungdahlii* strains with 27 mM fructose as the carbon source (150 mL headspace, 100 mL medium with culture). wt, *C. ljungdahlii* wt; Δ*aor1*, *C. ljungdahlii* Δ*aor1*; Δ*aor2, C. ljungdahlii* Δ*aor2*; Δ*aor1*Δ*aor2*, *C. ljungdahlii* Δ*aor1*Δ*aor2.* Lines represent the median of biological replicates (N=3) and triangles represent individual replicates (1 to 3). Numbers on growth rates, titers, and ratios are given in **Tab. S5**.

### 3.2 Ethanol titers are only impacted for *C. ljungdahlii* Δ*aor1* during growth with H_2_/CO_2_

Next, the growth of all strains was compared with H_2_ as the energy source and CO_2_ as the carbon source (**Fig. 2**). In contrast to growth with fructose, the OD_600_ was stable for all strains after reaching the maximum (**Fig. 2A)**. However, differences in the lag phases were observed between the strains (**Fig. 2A**). The pH dropped from 5.54 ± 0.02 to 3.80 ± 0.07 for all strains (**Fig. 2B**). While *C. ljungdahlii* wt consumed the provided CO_2_ completely, for all deletion strains, residual CO_2_ remained at the end of the experiment (**Fig. 2C**). The acetic acid production roughly followed the OD_600_ for all strains (**Fig. 2A,D**). Eventually, *C. ljungdahlii* Δ*aor1*Δ*aor2* reached an acetic acid titer that was comparable to *C. ljungdahlii* wt (*p*=0.930). However, both *C. ljungdahlii* Δ*aor1* and *C. ljungdahlii* Δ*aor2* reached significantly lower acetic acid titers (*p*=0.002 and *p*=0.016, respectively, **Fig. 2D, Tab. S5**). Significantly higher ethanol production was only achieved by *C. ljungdahlii* Δ*aor1* compared to *C. ljungdahlii* Δ*aor2* (p=0.007) and *C. ljungdahlii* wt (*p*=0.010, **Fig. 2E, Tab. S5**). Consequently, the highest E:A ratio using H_2_/CO_2_ was achieved by *C. ljungdahlii* Δ*aor1* despite being non-significant (*p*_kw_=0.189, **Fig. 2F, Tab. S5**). Previously, Xia et al. (2020) reported that the inactivation of *aor1* or *aor2* by the introduction of premature stop codons in *C. ljungdahlii* also led to decreased acetic acid titers for both strains with H_2_/CO_2_ compared to *C. ljungdahlii* wt. In contrast to our results here, ethanol production for the *aor1*-inactivated strain was lower than for *C. ljungdahlii* wt. However, it was also completely abolished for the *aor2*-inactivated strain (Xia et al., 2020). For *C. autoethanogenum*, only the *aor* double-deletion strain was investigated with H_2_/CO_2_ (Liew et al., 2017), which showed a similar growth pattern compared to *C. ljungdahlii* Δ*aor1*Δ*aor2* here (**Fig. 2**).

**Fig. 2.**
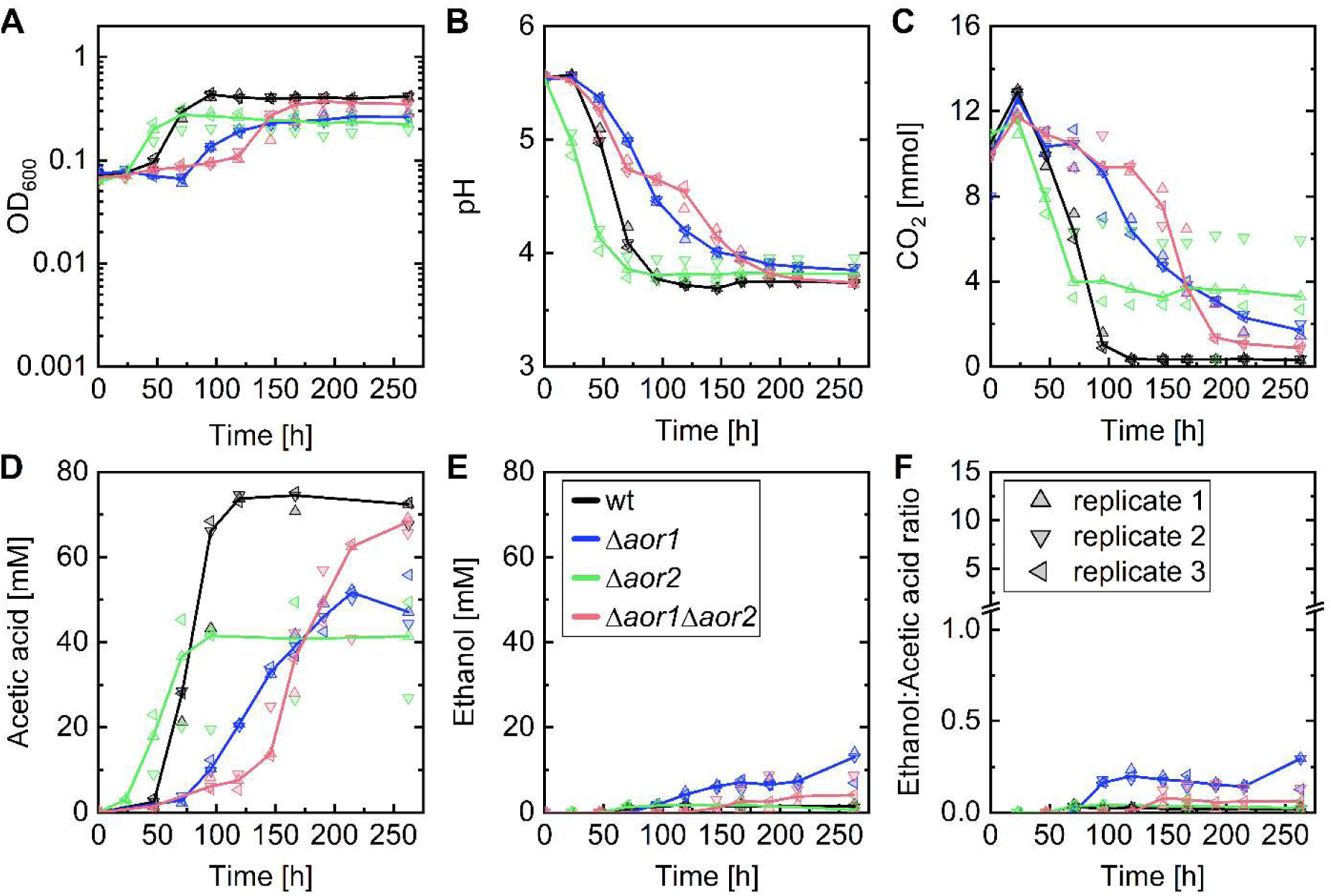
Bottle growth experiments with *C. ljungdahlii* strains using 1 bar overpressure of H_2_/CO_2_ as the carbon source (990 ml headspace, 100 ml medium with culture). wt, *C. ljungdahlii* wt; Δ*aor1*, *C. ljungdahlii* Δ*aor1*; Δ*aor2, C. ljungdahlii* Δ*aor2*; Δ*aor1*Δ*aor2*, *C. ljungdahlii* Δ*aor1*Δ*aor2.* Lines represent the median of biological replicates (N=3) and triangles individual replicates (1 to 3). Numbers on growth rates, titers, and ratios are given in **Tab. S5**.

### 3.3 Growth with CO shows the highest overall E:A ratios and reveals fluctuating ethanol production patterns

Lastly, all strains were compared for growth with CO (**Fig. 3**). We observed stochastic crash events, similar to previous reports with chemostats (Klask et al., 2020; Mahamkali et al., 2020). This did not allow us to identify statistically significant differences between the average values, and thus we discuss the results for individual replicates. All strains grew eventually but had long lag phases compared to growth on fructose or H_2_/CO_2_ (**Figs. 1A, 2A, 3A**). The pH dropped from ∼5.5 to ∼3.9 for *C. ljungdahlii* wt cultures but increased back to 5.1 for one of the replicates (**Fig. 3B**,▲). A similar behavior was observed for two replicates of *C. ljungdahlii* Δ*aor1* (**Fig. 3B**,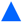,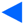). For these cultures, the pH decreased to 4.6 and 4.7 before it increased back to 5.4 and 5.8, respectively, followed by a second decrease of the pH (**Fig. 3B**,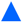,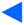). However, this behavior was not observed for *C. ljungdahlii* Δ*aor2* and *C. ljungdahlii* Δ*aor1*Δ*aor2* cultures (**Fig. 3B**). None of the cultures consumed the provided CO completely, but the CO consumption was highest for *C. ljungdahlii* wt (**Fig. 3C**). The highest acetic acid titers were reached for the *C. ljungdahlii* wt replicates that did not show the pH fluctuation (**Fig. 3D**,◄,▾**, Tab. S5**). In contrast, the *C. ljungdahlii* wt replicate with the pH rebound reached a much lower acetic acid titer, but the highest ethanol titer in the experiment (**Fig. 3E**,▲**, Tab. S5**). Thus, the E:A ratio for *C. ljungdahlii* wt for the replicate with pH fluctuation was high (**Fig. 3F**,▲**, Tab. S5**). This was also the case for the two replicates with pH fluctuations of *C. ljungdahlii* Δ*aor1* (**Fig. 3F**,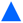,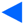**, Tab. S5**). *C. ljungdahlii* Δ*aor2* produced comparably high acetic acid levels, while ethanol titers remained low (**Fig. 3D,E, Tab. S5**). Overall, those E:A ratios with CO were the highest among the different carbon sources (**Figs. 1F, 2F, 3F, Tab. S5**). Similar or higher E:A ratios than those reported by us for *C. ljungdahlii* Δ*aor1* here, were only measured in chemostats and with additional feeding of acetic acid (Kwon et al., 2022; Richter et al., 2016; Schulz et al., 2023). To our knowledge, we are the first to describe an oscillating behavior in batch experiments with pure CO for *C. ljungdahlii*. Until now, oscillating patterns were described for chemostats with wt strains for: **1)** *C. autoethanogenum* fed with H_2_ and CO (Mahamkali et al., 2020); **2)** *C. ljungdahlii* fed with H_2_ and CO_2_ and an additional nitrate feed (Klask et al., 2020); and **3)** other *Clostridium* strains (Tyszak & Rehmann, 2024). All those studies discussed mechanisms that involved the depletion of metabolic intermediates and inhibitory effects on enzyme systems, leading to these oscillating patterns.

**Fig. 3.**
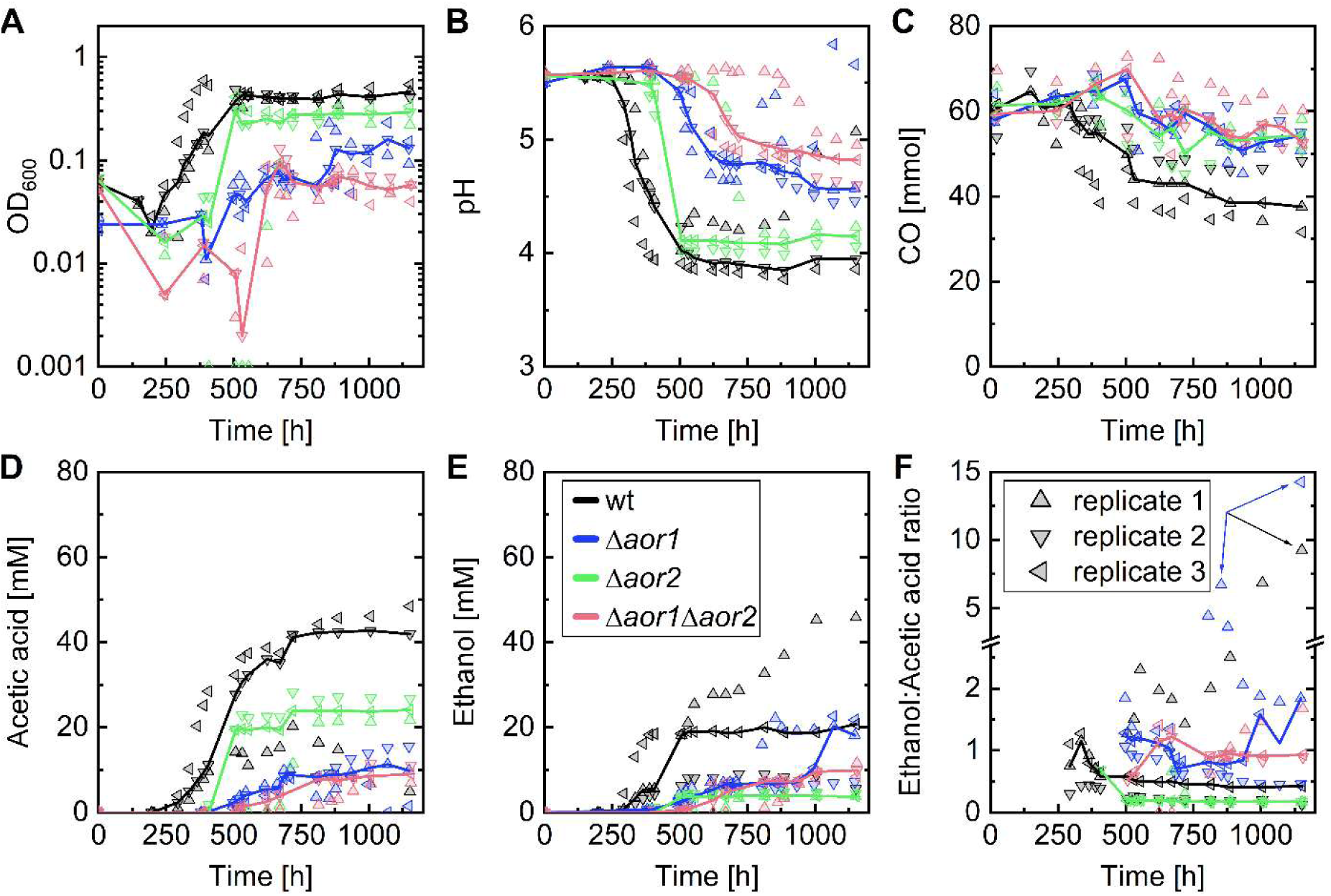
Bottle growth experiments with *C. ljungdahlii* strains using 1 bar overpressure of CO as the carbon source (990 ml headspace, 100 ml medium with culture). wt, *C. ljungdahlii* wt; Δ*aor1*, *C. ljungdahlii* Δ*aor1*; Δ*aor2, C. ljungdahlii* Δ*aor2*; Δ*aor1*Δ*aor2*, *C. ljungdahlii* Δ*aor1*Δ*aor2.* Lines represent the median of biological replicates (N=3) and triangles individual replicates (1 to 3). Blue and black arrows indicate the maximum E:A ratios for the individual bottles. Numbers on growth rates, titers, and ratios are given in **Tab. S5**.

Previously, it was shown that AORs are predominantly involved in ethanol production during autotrophic growth (indirect route), as opposed to the direct route, which is predominant during heterotrophic conditions (Richter et al., 2016; Tan et al., 2013; Whitham et al., 2015). Ethanol production was not completely abolished for the *C. ljungdahlii* Δ*aor1*Δ*aor2* strain with H_2_ and CO_2_ or with CO, and the E:A ratio was increased with fructose for this strain. Thus, there is, to some extent, an interplay between the direct and indirect route under all conditions. Furthermore, because the *C. ljungdahlii* Δ*aor1*Δ*aor2* strain still grew autotrophically, the microbe has a compensation method for losing the tungsten-dependent AORs. One option could be a switch to the direct route, while also the molybdenum-dependent AOR3 could play a role, although it was found only during oxygen-stress conditions before (Richter et al., 2016; Whitham et al., 2015).

Surprisingly, the shift in the product spectrum (*i.e.*, E:A ratio) that we measured for *C. ljungdahlii* Δ*aor1* and *C. ljungdahlii* Δ*aor2* were contrary to the effects observed for *C. autoethanogenum*, which might indicate regulatory differences in the two species (Liew et al., 2017). This suggests that AOR1 of *C. ljungdahlii* prefers converting acetaldehyde into acetic acid and AOR2 in the opposite direction, which rejects our hypothesis. Future research has to investigate the directionality of the AORs and the observed fluctuating patterns in more detail.

## Conclusions

We observed the highest ethanol production with CO as the carbon source. *C. ljungdahlii* Δ*aor1* achieved the highest E:A ratio of 14.3 with CO, but not in all replicates because we observed an oscillating pattern of pH, acetic acid, and ethanol production for *C. ljungdahlii* wt and *C. ljungdahlii* Δ*aor1* with CO. The experiments suggest an interplay of the direct and indirect route in ethanol production and different preferred catalytic directionalities for the AORs (AOR1 towards acetic acid and AOR2 towards acetaldehyde) because the E:A ratio trend for the deletion strains was the same for all carbon sources.

## Supplementary Material

Supplementary data of this work can be found in the online version of the paper.

## Research Data

Genome sequences of *C. ljungdahlii* wt, *C. ljungdahlii* Δ*aor1*, *C. ljungdahlii* Δ*aor2*, and *C. ljungdahlii* Δ*aor1*Δ*aor2* can be found in GenBank with the accession number PRJNA1118674.

## Acknowledgments

We thank Prof. Dr. Pengfei Xia for constructing the pMTLCas9 plasmid. We thank Dr. Andrés E. Ortiz Ardila for support with statistical analyses.

## Author contributions

Conceptualization: SS, STB, LTA, BM; Sequencing: JM, JAVG; Data curation: STB; Funding acquisition: SS, LTA, BM; Investigation: STB, SS; Methodology: STB, SS; Supervision: LTA, BM; Writing - original draft: STB, SS; Writing - review & editing: JM, JAVG, LTA, BM.

## Supplementary material

**Supplementary Table S1.** Comparison of wild type, *aor1*, and *aor2* deletion strains and their product spectrum in literature with various carbon sources. Given values are calculated from original data (Xia et al., 2020) or estimated from the figures and supplementary material (Liew et al., 2017). Fru, fructose; HAc, acetic acid; EtOH, ethanol; Ratio, ethanol:acetic acid ratio; n.c., not calculated; n.d., not determined.

|  |  | wild type |  |  | *aor1* deletion |  |  | *aor2* deletion |  |  | *aor1* and *aor2* deletion |  |  | Reference |
| --- | --- | --- | --- | --- | --- | --- | --- | --- | --- | --- | --- | --- | --- | --- |
|  |  | Fru | H<sub>2</sub>/CO<sub>2</sub> | CO | Fru | H<sub>2</sub>/CO<sub>2</sub> | CO | Fru | H<sub>2</sub>/CO<sub>2</sub> | CO | Fru | H<sub>2</sub>/CO<sub>2</sub> | CO |  |
| *C. ljungdahlii*<sup>a</sup> | HAc | n.d. | 62.29 ± 0.65 mM | n.d. | n.d. | 49.07 ± 0.12 mM | n.d. | n.d. | 40.13 ± 2.34 mM | n.d. | n.d. | n.d. | n.d. | Xia et al. (2020) |
|  | EtOH | n.d. | 1.86 ± 0.06 mM | n.d. | n.d. | 1.43 ± 0.01 mM | n.d. | n.d. | 0.00 ± 0.00 mM | n.d. | n.d. | n.d. | n.d. |  |
|  | Ratio |  | 0.03 |  |  | 0.03 |  |  | n.c. |  |  |  |  |  |
| *C. autoethanogenum* | HAc | 52 mM | 61 mM | 72 mM | 66 mM | n.d. | 66 mM | 52 mM | n.d. | 47 mM | 74 mM | 66 mM | 44 mM | Liew et al. (2017) |
|  | EtOH | 38 mM | 2 mM | 19 mM | 26 mM | n.d. | 8 mM | 58 mM | n.d. | 52 mM | 31 mM | 1 mM | 10 mM |  |
|  | Ratio | 0.73 | 0.03 | 0.26 | 0.39 |  | 0.12 | 1.11 |  | 1.11 | 0.42 | 0.02 | 0.23 |  |

<sup>a</sup>gene inactivation by the introduction of a premature stop codon *via* base editing

**Supplementary Table S2.**
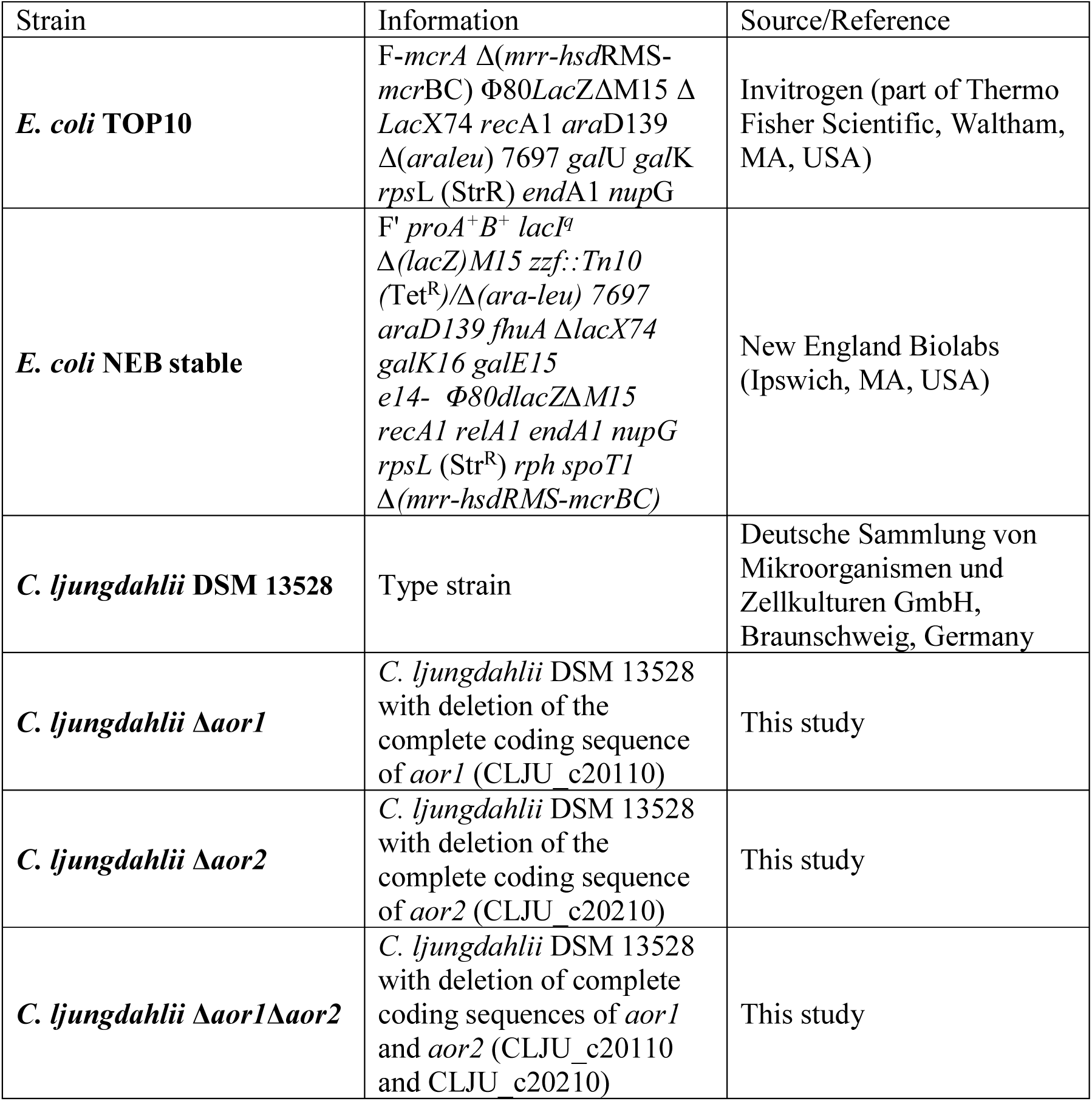
Bacterial strains used in this study.

**Supplementary Table S3.** Plasmids used in this study.

| Plasmid | Information | Source/Reference |
| --- | --- | --- |
| pMTLCas9 | *E. coli-Clostridium* shuttle vector based on pMTL84422 with anhydrotetracycline-inducible *cas9* | This study |
| pTargetF | *aadA*, P<sub>J23119</sub> for constitutive sgRNA cassette expression | Jiang et al. (2015) |
| pgRNA_*aor1* | pTargetF containing sgRNA cassette targeting *aor1* (CLJU_c20110) | This study |
| pgRNA_*aor2* | pTargetF containing sgRNA cassette targeting *aor2* (CLJU_c20210) | This study |
| pMTLCas9_*aor1* | pMTLCas9 containing sgRNA cassette targeting *aor1* (CLJU_c20110) and homologous regions for deletion of *aor1* | This study |
| pMTLCas9-*aor2*-STEP1 | pMTLCas9 containing sgRNA cassette targeting *aor2* (CLJU_c20210) | This study |
| pMiniT2.0 | *bla*, vector for cloning | New England Biolabs (Ipswich, MA, USA) |
| pMiniT2.0_donor-*aor2* | pMiniT2.0 with homologous regions for deletion of *aor2* (CLJU_c20210) | This study |
| pMTLCas9_*aor2* | pMTLCas9 containing sgRNA cassette targeting *aor2* (CLJU_c20210) and homologous regions for deletion of *aor2* | This study |
| pANA1 | *bla*, Φ3T I methyl transferase for methylation of plasmids | Molitor et al. (2016) |

**Supplementary Table S4.**
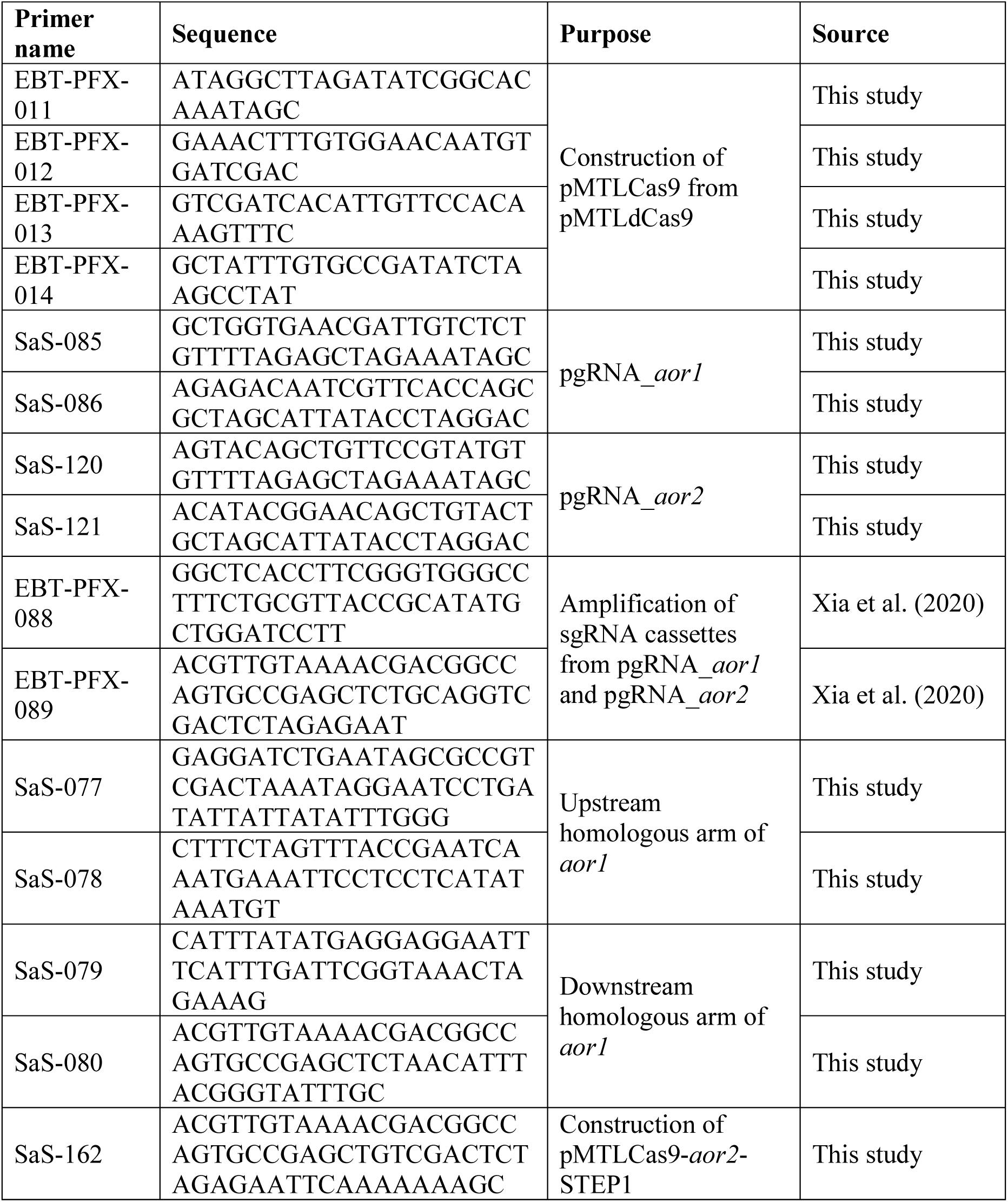
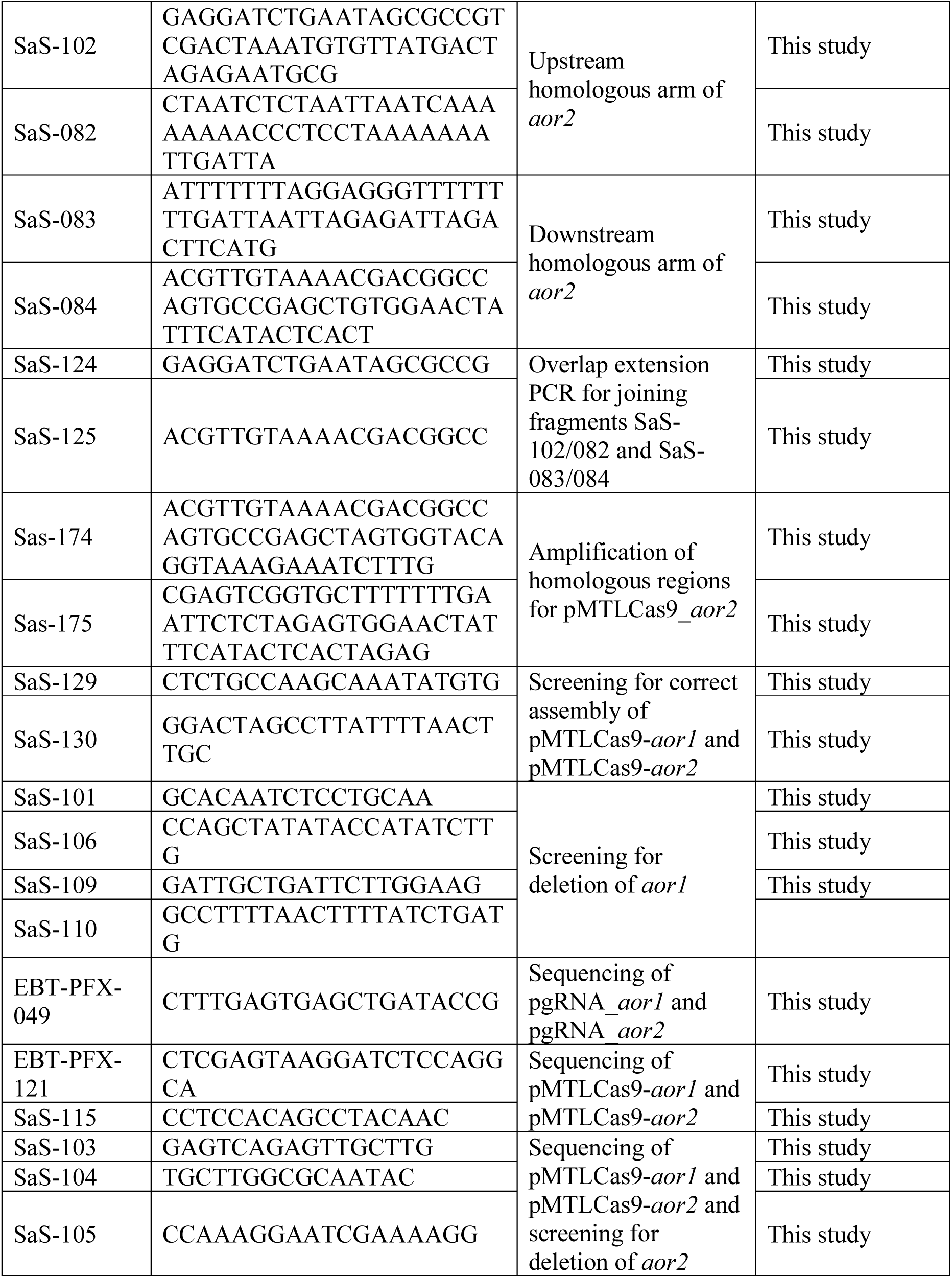
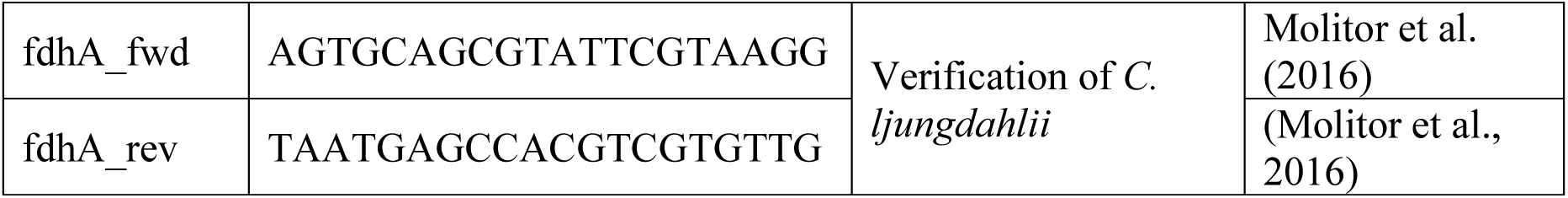
Primers used in this study.

**Supplementary Table S5.**
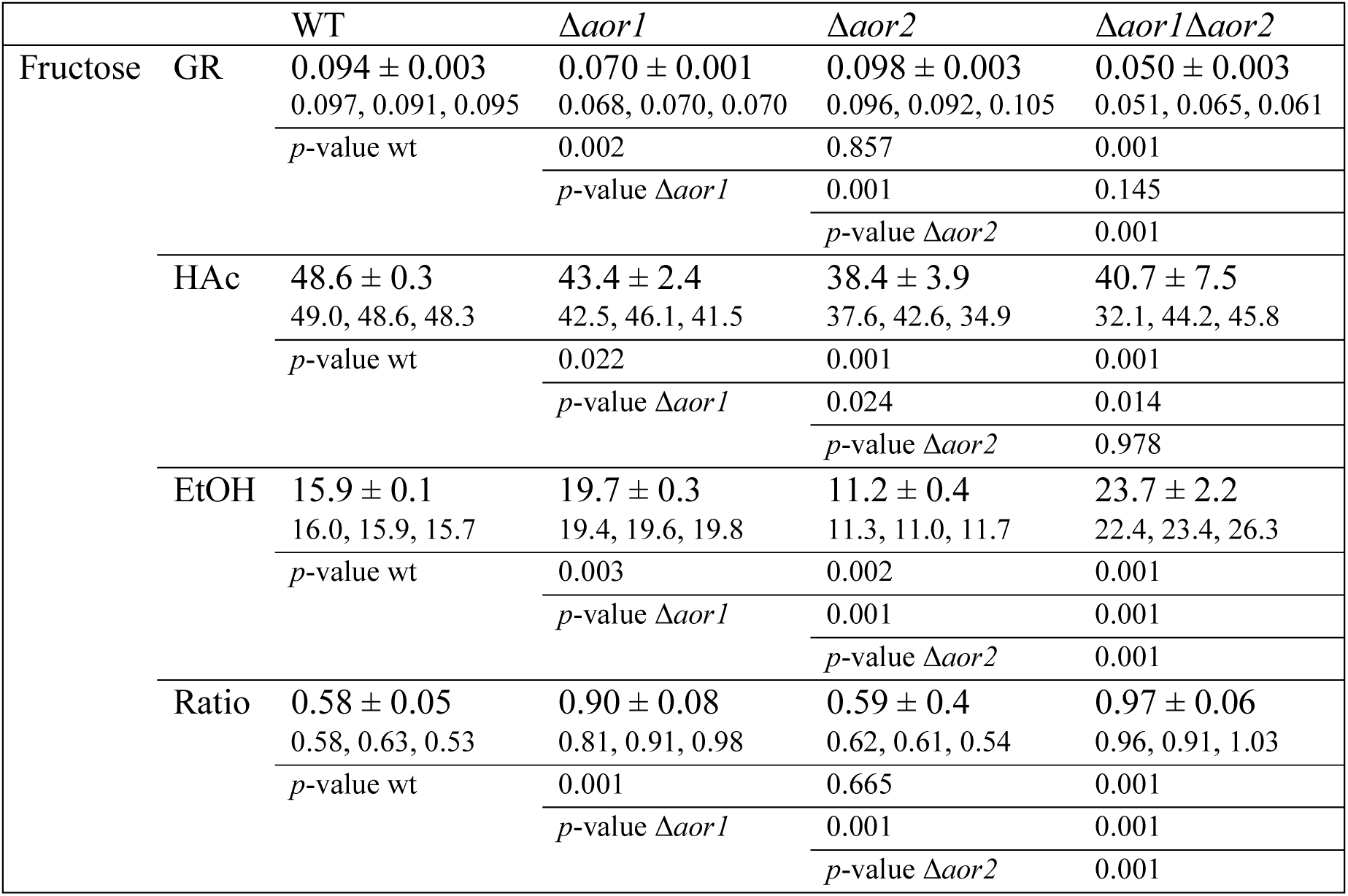
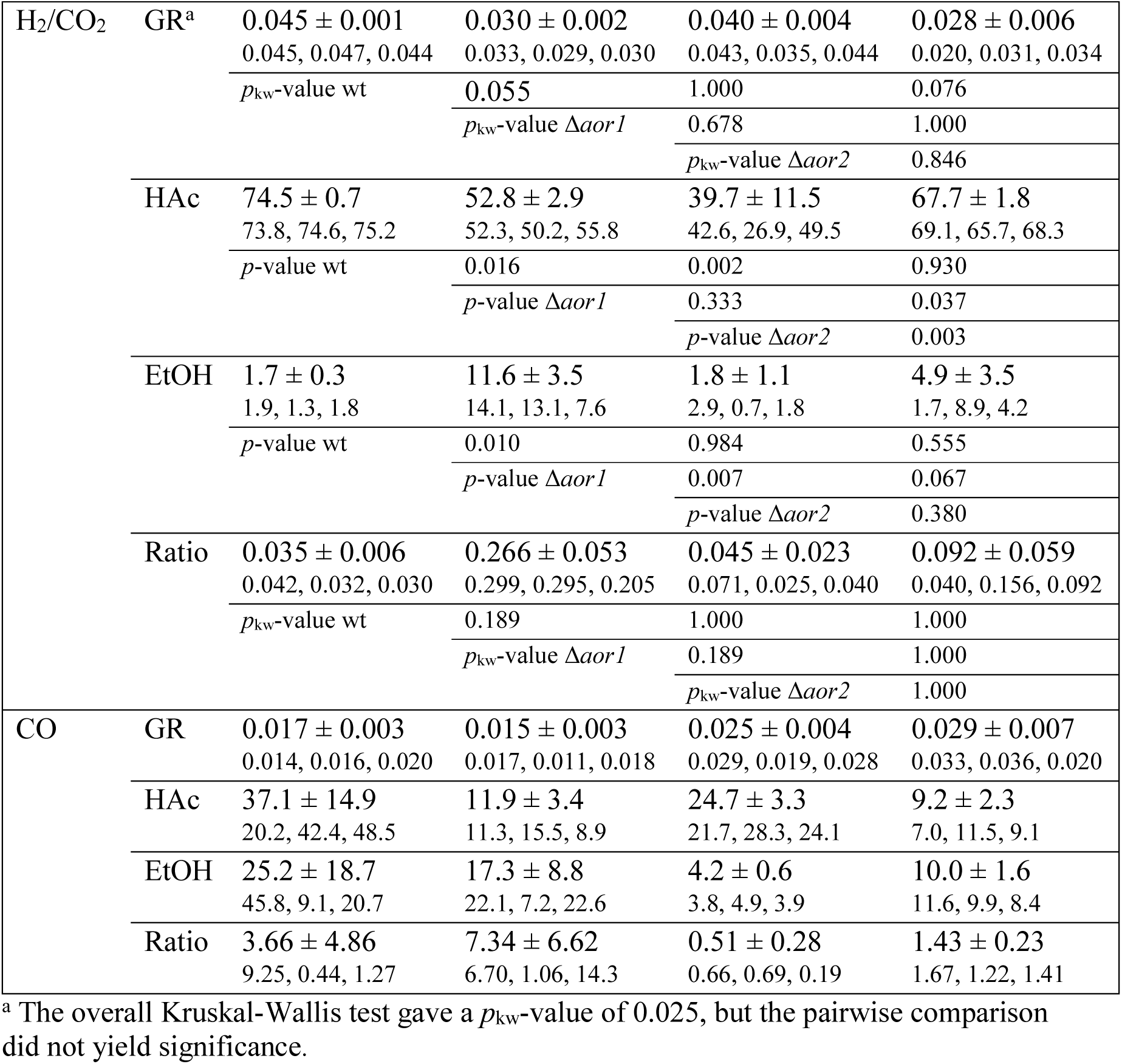
Maximum growth rate (GR, in h^-1^), titers of acetic acid (HAc) and ethanol (EtOH) (in mmol L^-1^), and ethanol:acetic acid ratios (Ratio) achieved by the different *C. ljungdahlii* strains. *C. ljungdahlii* wild type (WT), *C. ljungdahlii* Δ*aor1* (Δ*aor1*), *C. ljungdahlii* Δ*aor2* (Δ*aor2*), and *C. ljungdahlii* Δ*aor1*Δ*aor2* (Δ*aor1*Δ*aor2*) grown with different substrates (*i.e.*, fructose, H_2_/CO_2_ [80:20 vol-%], or CO). The ratios were calculated from acetic acid and ethanol titers throughout the growth experiments. Means ± standard deviation were calculated from triplicates with the maximum titers and numbers in the line below. *p*-values are given in comparison to the indicated strains in the first column of the respective line (*p*, ANOVA-based; *p*_kw_, Kruskal-Wallis-based because criteria for the parametric test were not met). No statistics were calculated for the growth experiment using CO as carbon source.

